# Meta-analytic evidence for the plurality of mechanisms in transdiagnostic structural MRI studies of hallucination status

**DOI:** 10.1101/413609

**Authors:** Colleen Rollins, Jane R Garrison, Jon S Simons, James B Rowe, Claire O’Callaghan, Graham Murray, John Suckling

**Author notes:** Correspondence to: Ms. Colleen Rollins, Department of Psychiatry, University of Cambridge, Cambridge CB2 0SP, UK, 07511 537162.

## Abstract

**BACKGROUND:** Hallucinations are transmodal and transdiagnostic phenomena, occurring across sensory modalities and presenting in psychiatric, neurodegenerative, neurological, and non-clinical populations. Despite their cross-category occurrence, little empirical work has directly compared between-group neural correlates of hallucinations.

**METHODS:** We performed whole-brain voxelwise meta-analyses of hallucination status across diagnoses using AES-SDM, and conducted a comprehensive systematic review in PubMed and Web of Science until May 2018 on other structural correlates of hallucinations, including cortical thickness and gyrification.

**FINDINGS:** 3214 abstracts were identified. Patients with psychiatric disorders and hallucinations (eight studies) exhibited reduced gray matter (GM) in the left insula, right inferior frontal gyrus, left anterior cingulate/paracingulate gyrus, left middle temporal gyrus, and increased in the bilateral fusiform gyrus, while patients with neurodegenerative disorders with hallucinations (eight studies) showed GM decreases in the left lingual gyrus, right supramarginal gyrus/parietal operculum, left parahippocampal gyrus, left fusiform gyrus, right thalamus, and right lateral occipital gyrus. Group differences between meta-analyses were formally confirmed and a jackknife sensitivity analysis established the reproducibility of results across nearly all study combinations. For other measures (28 studies), the most consistent findings associated with hallucination status were reduced cortical thickness in temporal gyri in schizophrenia and altered hippocampal volume in Parkinson’s disease and dementia.

**INTERPRETATION:** Distinct patterns of neuroanatomical alteration characterize hallucination status in patients with psychiatric and neurodegenerative diseases, suggesting a plurality of anatomical signatures. This approach has implications for treatment, theoretical frameworks, and generates refutable predictions for hallucinations in other diseases and their occurrence within the general population.

**FUNDING:** None.

**Research in context:** *Evidence before this study:* There is increasing recognition that hallucinations occur beyond the archetype of schizophrenia, presenting in other psychiatric disorders, neurological and neurodegenerative conditions, and among the general population. Not only are hallucinations a transdiagnostic phenomenon, but also the experience of hallucinating is phenomenologically diverse, varying in modality, content, frequency, and affect. It has been suggested that no one type of hallucination is pathognomic to any one disorder, but rather that hallucinations may exist on a continuum. However, limited research has been done to directly compare the underlying neuroanatomy of hallucinations between different disorders. With this aim, we conducted a meta-analysis and systematic review of structural MRI studies comparing individuals who experience hallucinations with those who do not, to investigate the brain morphology related to the transdiagnostic presentation of hallucinations. We searched PubMed and Web of Science with no start date limit, up to May 2018 using the keyword combination (hallucinat*) AND (MRI OR magnetic resonance imaging OR morphology OR voxel?based OR morphometr* OR neural correlate OR structur*). We included only studies with a within-group no-hallucination control to tease out structural changes specific to hallucinations from effects of the broader pathology. Neuroimaging meta-analyses were conducted on studies performing whole-brain voxelwise gray matter differences, while studies assessing other structural correlates were qualitatively synthesized.

*Added value of this study:* This is the first meta-analysis to illustrate the brain structural correlates of hallucination occurrence derived from T1-weighted MRI, and to do so in a comparative manner across clinical groups. We identified two distinct gray matter substrates for hallucination presence in psychiatric compared to neurodegenerative diseases, which we hypothesise constitute at least two distinct mechanisms. In addition, we qualitatively assessed other structural neuroimaging studies over a variety of morphometric indices. We therefore provide a complete characterization of current knowledge of the brain morphology associated with hallucinations across clinical status and modality.

*Implications of all the available evidence:* Our findings show at least two structural substrates that link to the hallucinatory experience. This informs theoretical work on hallucinations which have to date been limited in generating unifying direction-specific predictions of brain structure and function. Understanding the plurality of anatomical signatures of hallucinations may also inform treatment strategies. We predict that other disorders in which patients experience hallucinations can be categorised by our approach based on the broader phenotype; for example, hallucinations in personality disorder may be of the psychiatric type, and similarly for early onset hallucinations in the general population, whilst later onset will be neurodegenerative. Moreover, by differentiating the mechanisms of hallucinations we recommend the contextualising of research by the appropriate phenotype.

## Introduction

Hallucinations are transdiagnostic and transmodal perceptions of stimuli that do not exist in the physical world^1^. They are prevalent in both psychiatric disorders, such as schizophrenia (60-80%)^2^ and bipolar disorder (BD 10-23%)^3^, and neurodegenerative diseases, such as Parkinson’s disease (PD; 22-38%)^4^, dementia with Lewy Bodies (DLB; 80%)^5^, and Alzheimer’s disease (AD; 13-18%)^6^, as well as in other psychiatric and neurological disorders, and among the general population (4.5-12.7%)^7^. Irrespective of diagnosis, the presence of hallucinations marks an increased risk of adverse outcomes, such as reduced likelihood of recovery in schizophrenia^8^, more severe cognitive deficits in PD^9^, increased mortality in AD^10^, increased suicidal behaviour in adults with psychosis^11^, and transition to later mental illness in children and young adults^12,13^. Although hallucinations are often distressing, they may also be benign or contribute to meaningful personal experiences^14,15^.

Historically, hallucinations were considered a cardinal symptom of schizophrenia, but they are not pathognomic: one-third of patients do not hallucinate^2^, and the experience is often heterogeneous among those who do^1^. This has been confirmed across clinical and non-clinical populations, revealing diverse phenomenology involving modality, content, affect, onset, and frequency^15,16^. Inter-individual differences among hallucinations prompt a number of conceptual, mechanistic, and clinical questions: Does phenomenological heterogeneity translate into neurobiological plurality? How would this influence theoretical models of hallucinations and inform treatments? Does the epidemiological and experiential diversity of hallucinations reflect a continuum model, in which symptoms like hallucinations are distributed over a spectrum of individuals who do and do not meet criteria for mental illness, and thus arise from a common mechanism instantiated to different degrees of severity^17^? Establishing the validity of this conceptual framework against alternatives is important for how we understand and treat hallucinations.

Despite the plurality of hallucinations, there is little empirical work comparing between-group neural correlates of hallucinations. Prior reviews and meta-analyses on the brain structural and functional correlates of hallucinations have generally limited their scope to a single diagnosis or modality^18-20^, or both^21-25^. Only two reviews have investigated hallucinations transdiagnostically or in more than one modality: one without quantitative meta-analytic comparison^26^, the other focussed on acute functional correlates of hallucinations^27^. Two meta-analyses have explored the structural correlates of hallucinations, but assessed correlates of hallucination severity rather than presence/absence, and limited their scope to auditory verbal hallucinations (AVH) in schizophrenia^23,24^. We therefore planned meta-analyses to evaluate MRI-derived volumetric structural grey matter (GM) correlates of hallucination status across populations, complemented with a comprehensive review of other structural measures, including cortical thickness, gyrification, and structure-specific morphometrics.

A significant issue in neuroimaging studies of hallucinations has been the lack of a clinical control group, thus confounding abnormalities specific to hallucination status with those of the broader phenotype. Equally challenging has been a tangled conceptual landscape, with numerous models proposed as cognitive or neurobiological accounts of auditory or visual hallucinations^5,26,28-41^ (Figure 1). Obtaining differentiating evidence is difficult as these models are not mutually exclusive, each drawing upon a similar repertoire of constituents, making it non-trivial to derive corresponding predictions^42^. However, specific morphological variation can differentiate patients who do and do not hallucinate^43^, indicating that structural MRI can provide insights into why individuals hallucinate.

**Figure 1.**
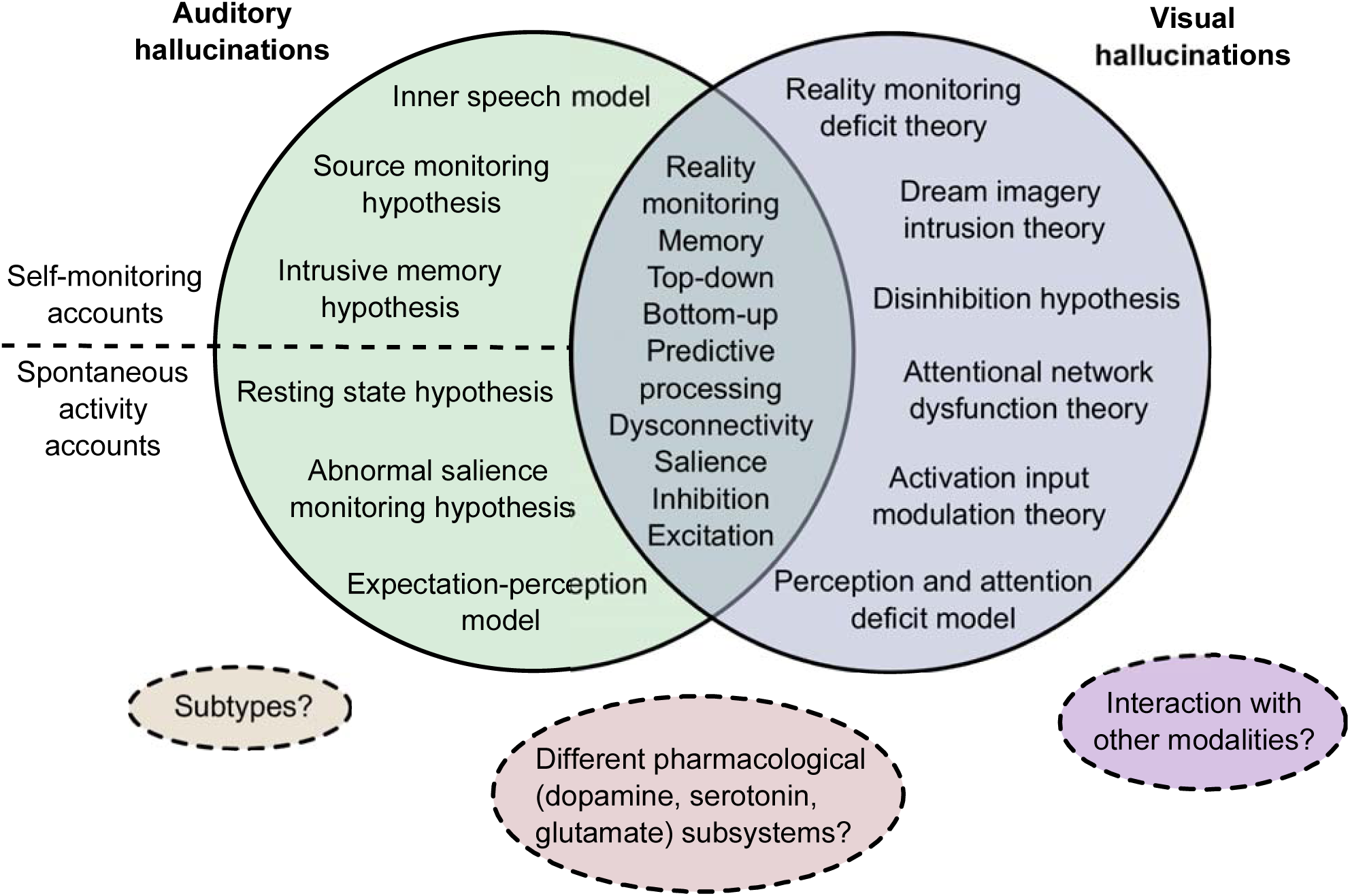
Landscape of theoretical models of hallucinations. The major cognitive, psychological, and neurobiological theories for auditory and visual hallucinations are depicted. Separate theories have been proposed to underlie auditory versus visual hallucinations, although they share many common themes. Different theories within each modality category are not mutually exclusive and may overlap in their predictions. Dotted lines delineate proposals of divisions between, extensions to, or limitations of current theories. **Key references:** Inner speech model^45^; Intrusive memory hypothesis^28^; Resting state hypothesis^29^; Abnormal salience monitoring hypothesis^30^; Expectation-perception model^31^; Reality monitoring deficit theory, Dream imagery intrusion theory, Activation input modulation theory^36,46^; Disinhibition hypothesis^5^; Perception and attention deficit model^32^; Top-down bottom-up models^26,33^; Excitatory-inhibitory imbalance^34^; Predictive processing accounts^35,37^; proposal of divide between self-monitoring accounts and spontaneous activity accounts for auditory verbal hallucinations (AVH)^38^; proposal of subtypes for AVH^40^; proposal for differential contribution of pharmacological subsystems to different types of AVH^41^; commentary on need to address interaction between and hierarchy of different modalities of hallucinations^109^.

Voxel-based morphometry (VBM) is a common method for unbiased, automated quantification of GM differences between groups. Conducting a meta-analysis of VBM studies is an objective approach to synthesize the extant literature and identify replicable findings^44^. Knowledge of neuroanatomical signatures of hallucinations present in certain populations and absent in others would clarify the continuum model and contribute towards a clearer neurobiological picture of the origins and mechanisms of hallucinations. Considering the cultural and historical influences on hallucination interpretation^14^, an organic model of hallucinations could moreover substantiate accurate diagnostic criteria. This meta-analysis and systematic review quantitatively compared people with and without hallucinations in terms of brain structure to identify the neuroanatomy related to the transdiagnostic presence of hallucinations.

## Methods

### Search strategy and selection criteria

A systematic review of the literature for the structural correlates of hallucinations was conducted in October 2017, with update notifications received until May 2018. Following PRISMA guidelines^47^, articles were identified by searching PubMed and Web of Science using the keyword combination (hallucinat*) AND (MRI OR magnetic resonance imaging OR morphology OR voxel?based OR morphometr* OR neural correlate OR structur*) with no date limit. Reviews and meta-analyses on neuroimaging of hallucinations were cross-referenced to ensure no relevant studies were missed.

Studies were included if they: (a) employed structural MRI in a whole-brain investigation of voxelwise differences in GM reported in standard stereotaxic space; (b) included a direct comparison between groups with and without hallucinations within the same diagnostic category. Corresponding authors were contacted to request coordinate information if not reported in the original article, or to clarify methodological issues. CR evaluated all studies and JS, GM, or JRG confirmed the selection criteria, with uncertainties discussed to consensus. Region of interest (ROI) VBM studies and studies using non-voxelwise structural MRI methods that otherwise matched inclusion criterion (b) were included in the systematic review.

### Data analysis

Voxel-wise meta-analyses were undertaken using anisotropic effect-size seed-based d Mapping (AES-SDM; https://www.sdmproject.com/)^48,49^ following recommended guidelines ^44^ (Supplementary S1). AES-SDM uses peak coordinates and effect sizes from primary studies to create maps of meta-analytic effect size and variance of the signed GM differences. Similar to other voxel-based meta-analytic methods^50^, loci from primary studies are estimated as smoothed spheres and meta-analytic maxima calculated by weighting the encompassed voxels^48^. Additionally, AES-SDM incorporates the effect sign (increases or decreases) and the t-statistic associated with each peak, increasing both sensitivity and accuracy^48^. AES-SDM also allows inclusion of non-significant studies, reducing bias towards positive results. AES-SDM is detailed elsewhere (https://www.sdmproject.com/software/tutorial.pdf), and summarized in Supplementary S2.

Anticipating differences in mechanisms of hallucinations between psychiatric illnesses and neurodegenerative diseases based on distinctions in phenomenology, modality, prevalence^51^, and the significant participant age separation amongst primary studies (t(25) = 17.324, p<0.001), we performed a meta-analysis including schizophrenia, first episode schizophrenia (FES), first episode psychosis (FEP), and young adults at clinical risk for psychosis (at-risk mental state long-term, ARMS-LT), and BD, and a second of neurodegenerative disorders, including PD and AD. Of the 16 studies included in these two cross-sectional meta-analyses, three (see Table 1) did not make an explicit comparison between a hallucination (H) and no-hallucinations (NH) group, though the majority of patients in each group respectively either did or did not have hallucinations, and were therefore included^52-54^. A jackknife sensitivity analysis was performed on the meta-analyses to test reproducibility of significant brain regions by iteratively repeating the statistical analysis systematically excluding one study^55^. Finally, we formally assessed group differences between psychiatric and neurodegenerative hallucination meta-analyses using Monte Carlo randomizations to determine statistical significance^56^.

**Table 1.** Demographic and clinical characteristics of included studies.

| Group | Study | Sample | N | Age (SD) | M/F | Hallucination Assessment Scale | Modality |
| --- | --- | --- | --- | --- | --- | --- | --- |
| Psychiatric | Garrison et al., 2015 [43] | SCZ-H<br>SCZ-NH | 79<br>34 | 38.5 (9.8)<br>40.7 (9.8) | 65/14<br>27/7 | clinical interview | mixed |
|  | Gaser et al., 2004 [57] | SCZ-H<br>SCZ-NH | 29<br>56 | 36.2 (10.9)<sup>†</sup> | 52/33<sup>†</sup> | SAPS | auditory |
|  | Shapleske et al., 2002 [58] | SCZ-H<br>SCZ-NH | 41<br>31 | 35.5 (8.8)<br>32.0 (7.5) | 41<br>31 | SAPS | auditory |
|  | van Swam et al., 2012<sup>c</sup> [52] | SCZ-H<br>SCZ-NH | 10<br>10 | 40.9 (8.8)<br>36.3 (5.6) | 5/5<br>7/3 | PANSS, semi-structured interview | auditory |
|  | van Tol et al., 2014 [59] | SCZ-H<br>SCZ-NH | 31<br>20 | 33.4 (12.5)<br>35.0 (9.7) | 27/4<br>17/3 | PANSS | auditory |
|  | Huang et al., 2015 [60] | FES-H<br>FES-NH | 18<br>18 | 22.6 (6.7)<br>22.7 (3.9) | 10/8<br>9/9 | PANSS, HAHRS | auditory |
|  | Smieskova et al., 2012<sup>ab</sup> [53] | FEP-H<br>ARMS-LT-NH | 16<br>13 | 25.1 (4.6)<br>24.6 (2.2) | 12/4<br>8/5 | BRPS | auditory |
|  | Neves et al., 2016 [61] | BD-H<br>BD-NH | 9<br>12 | 37.7 (12.1)<br>39.9 (15.0) | 3/6<br>6/6 | MINI-Plus | auditory or visual |
| Neuro-degenerative | Goldman et al., 2014 [62] | PD-H<br>PD-NH | 25<br>25 | 74.8 (6.0)<br>75.4 (6.1) | 17/8<br>18/7 | MDS-UPDRS | mixed |
|  | Meppelink et al., 2011 [63] | PD-H<br>PD-NH | 11<br>13 | Not reported | Not reported | NPI | visual |
|  | Pagonbarraga et al., 2014 [64] | PD-H<br>PD-NH | 15<br>27 | 64.1 (9)<br>66.3 (8) | Not reported | MDS-UPDRS | passage and/or presence |
|  | Ramirez-Ruiz et al., 2007 [65] | PD-H<br>PD-NH | 18<br>20 | Not reported | 8/12<br>7/11 | NPI (Spanish version), semi-structured interview | visual |
|  | Watanabe et al., 2013 [66] | PD-H<br>PD-NH | 13<br>13 | 66.6 (5.5)<br>63.6 (10.7) | 7/6<br>5/8 | UPDRS | visual |
|  | Shin et al., 2012 [67] | nPD-H<br>nPD-NH | 46<br>64 | 71.3 (5.9)<br>70.7 (5.7) | 26/38<br>18/9 | NPI | visual |
|  | Lee et al., 2016<sup>a</sup> [54] | AD-H<br>AD-NH | 17<br>25 | 74.3 (7.3)<br>72.4 (9.4) | 4/13<br>6/19 | NPI (Korean version) | auditory or visual |
|  | Blanc et al., 2014 [68] | AD-H<br>AD-NH | 39<br>39 | 76.0 (7.4)<br>76.4 (7.2) | 20/19<br>20/19 | NPI | auditory or visual |

Abbreviations: AD: Alzheimer's disease; PD: Parkinson's disease; SCZ: schizophrenia; FES: first episode schizophrenia; BD: bipolar disorder; nPD: Parkinson's disease without dementia; FEP: first episode psychosis; ARMS-LT: at risk mental state long-term; X-H: population X with hallucinations; X-NH: population X without hallucinations; NPI: Neuropsychiatric Inventory Questionnaire; MDS-UPDRS: Movement Disorder Society (MDS)-sponsored version of the Unified Parkinson's disease Rating Scale (UPDRS); PANSS: Positive and Negative Symptom Scale; HAHRS: Hoffman Auditory Hallucination Rating Scale; MINI-Plus; Mini International Neuropsychiatric Interview (MINI) Plus; SAPS: Scale for the Assessment of Positive Symptoms; BRPS: Brief Psychiatric Rating Scale. Dashed
\* Studies considered proxy comparisons between hallucinating and non-hallucinating groups.
<sup>a</sup> Lee et al. (2016) compared AD patients with misidentification subtype to AD patients without psychosis, though they classified AD patients with hallucination into the misidentification subtype.
<sup>b</sup> Smieskova et al. (2012) compared FEP to ARMS-LT participants, though the groups differed significantly ( $p < 0.0001$ ) in their hallucination score, with the FEP group having a mean (S.D.) score of 3.5 (2.0) on the BRPS hallucination item 10 (moderate – moderately severe) and the ARMS-LT group having a mean score of 1.4 (1.0) – a score of 1 being the lowest possible score.
<sup>c</sup> Van Swam et al. (2012) used voxel-wise cortical thickness, as opposed to VBM. Though a different analysis, VWCT and VBM are considered complementary methods<sup>95</sup>.
<sup>†</sup> For total sample of patients with schizophrenia, including both H and NH.

### Role of the funding source

There was no funding source for this study. CR had full access to all the data in the study and had final responsibility for the decision to submit for publication.

## Results

The literature search identified 2259 articles from PubMed and 1785 from Web of Science, for a merged total of 3214 after duplicates were excluded (Figure 2). 99 articles were selected for whole text retrieval after title/abstract screening. 16 studies met criteria for the meta-analyses^43,52-54,57-68^ (see Table 1 for sample characteristics; Table 2 for analysis details and results summary) and 28 papers (18 psychiatric; 10 neurodegenerative) for the systematic review of other structural metrics comparing groups with and without hallucinations.

**Table 2.** Imaging characteristics and key results of included studies.

| Group | Study | T | Software | Covariates | FWHM (mm) | Statistical Threshold | Original stereotaxic space | n Foci | Main result |
| --- | --- | --- | --- | --- | --- | --- | --- | --- | --- |
| Psychiatric | Garrison et al., 2015 [43] | 1.5 | SPM8 | age, gender | 8 | p<0.001, uncorrected; minimum cluster size=100 voxels | MNI | 2 | H>NH: bilateral occipital lobe |
|  | Gaser et al., 2004 [57] | 1.5 | SPM99 | SANS total score, SAPS total score without auditory hallucination sub-items, gender | 8 | p<0.001, uncorrected, k=100 voxels | Talraich | 4 | H<NH: L transverse temporal (Heschl's) gyrus R middle/inferior frontal gyrus, L middle temporal gyrus, L paracingulate gyrus, |
|  | Shapleske et al., 2002 [58] | 1.5 | AFNI | age and handedness | -4.2 | absolute value of standard error <1.96 | Talairach | 1 | H<NH: L insular cortex |
|  | van Swam et al., 2012 [52] | 3 | Brain Voyager QX 1.9 | none | Not reported | p<0.05, cluster size<15 voxels, corrected for multiple comparisons (Bonferroni p < 0.0063) | MNI | 7 | H>NH: L middle frontal gyrus, L posterior cingulate gyrus, L frontal insula, L parahippocampal gyrus, L postcentral sulcus, R visual cortex<br>H<NH: posterior inferior temporal sulcus, postcentral gyrus |
|  | van Tol et al., 2014 [59] | 3 | SPM8 | age, sex | 8 | p<0.05 FWE-corrected (cluster level), voxel-wise threshold of p<0.005 uncorrected | Talairach | 3 | H<NH: L putamen |
|  | Huang et al., 2015 [60] | 3 | SPM8 | age, gender, years of education | 8 | p<0.001, uncorrected | Talairach | 0 | n.s. |
|  | Smieskova et al., 2012 [53] | 3 | SPM8 | age, gender and total GMV | 8 | p<0.001, uncorrected (cluster-forming threshold); p<0.05 FWE-corrected | Talairach | 5 | H<NH: L parahippocampal gyrus<br>H>NH: L superior frontal gyrus, L caudate |
|  | Neves et al., 2016 [61] | 1.5 | SPM8 | total GMV | 8 | p<0.05, corrected for FWE across the entire brain | Not reported | 0 | n.s. |
| Neuro-degenerative | Goldman et al., 2014 [62] | 1.5 | SPM8 | total intracranial volume | 8 | p<0.01, uncorrected; cluster extent threshold k=10 | MNI | 18 | H<NH: bilateral cuneus, bilateral fusiform gyrus, bilateral inferior parietal lobule, bilateral precentral gyrus, bilateral middle occipital gyrus, R lingual gyrus, bilateral cingulate gyrus, L paracentral lobule |
|  | Meppelink et al., 2011 [63] | 3 | SPM5 | total GM | 10 | p<0.05, brain-volume corrected cluster-level | MNI | 0 | n.s. |
|  | Pagonbarraga et al., 2014 [64] | 1.5 | SPM5 | age, gender and global GMV | 12 | p<0.001, uncorrected; cluster size=207 voxels (determined by 1000 Monte Carlo simulations) | Not reported | 4 | H<NH: R vermis, R precuneus<br>H>NH: posterior lobe of cerebellum, L inf. frontal cortex |
|  | Ramirez-Ruiz et al., 2007 [65] | 1.5 | SPM2 | TIV, MMSE, Hamilton and Hoehn and Yahr scores | 12 | p<0.05, corrected cluster p-level | MNI | 3 | H<NH: bilateral sup. parietal lobe, L lingual gyrus |
|  | Watanabe et al., 2013 [66] | 3 | SPM8 | total intracranial volume, age, and sex | 8 | p<0.01, FWE corrected; cluster size > 50 voxels and z-scores > 3.00 | MNI | 15 | H<NH: bilateral middle frontal gyrus, L cingulate gyrus, R inferior parietal lobule, bilateral cuneus, L fusiform gyrus, L posterior lobe, L inferior occipital gyrus, L inferior frontal gyrus, L declive, R lingual gyrus |
|  | Shin et al., 2012 [67] | 3 | SPM8 | age, sex, PD duration, intracerebral volume, K-MMSE score | 6 | p<0.05, FWE corrected, and at a more liberal threshold of uncorrected p<0.001 at the voxel level with a minimum cluster size of 100 voxel | Talairach | 5 | H<NH: R inferior frontal gyrus, L thalamus, L uncus, L parahippocampal gyrus |
|  | Lee et al., 2016 [54] | 3 | SPM8 | age, gender, education, CDR score, NPI non-psychotic scores | 8 | p<0.001 uncorrected; extent threshold of contiguous 100 voxels (k>100) | Not reported | 6 | H<NH: R inferior parietal lobule, R lingual gyrus, L cuneus, R middle frontal gyrus, R superior occipital gyrus, R middle temporal gyrus |
|  | Blanc et al., 2014 [68] | 1.5 | SPM12b | age, total amount of GM | 8 | p<0.001, uncorrected; minimum cluster size=25 voxels | Not reported | 3 | H<NH: R insula / inferior frontal gyrus, L superior frontal gyrus, bilateral lingual gyrus |

**Figure 2.**
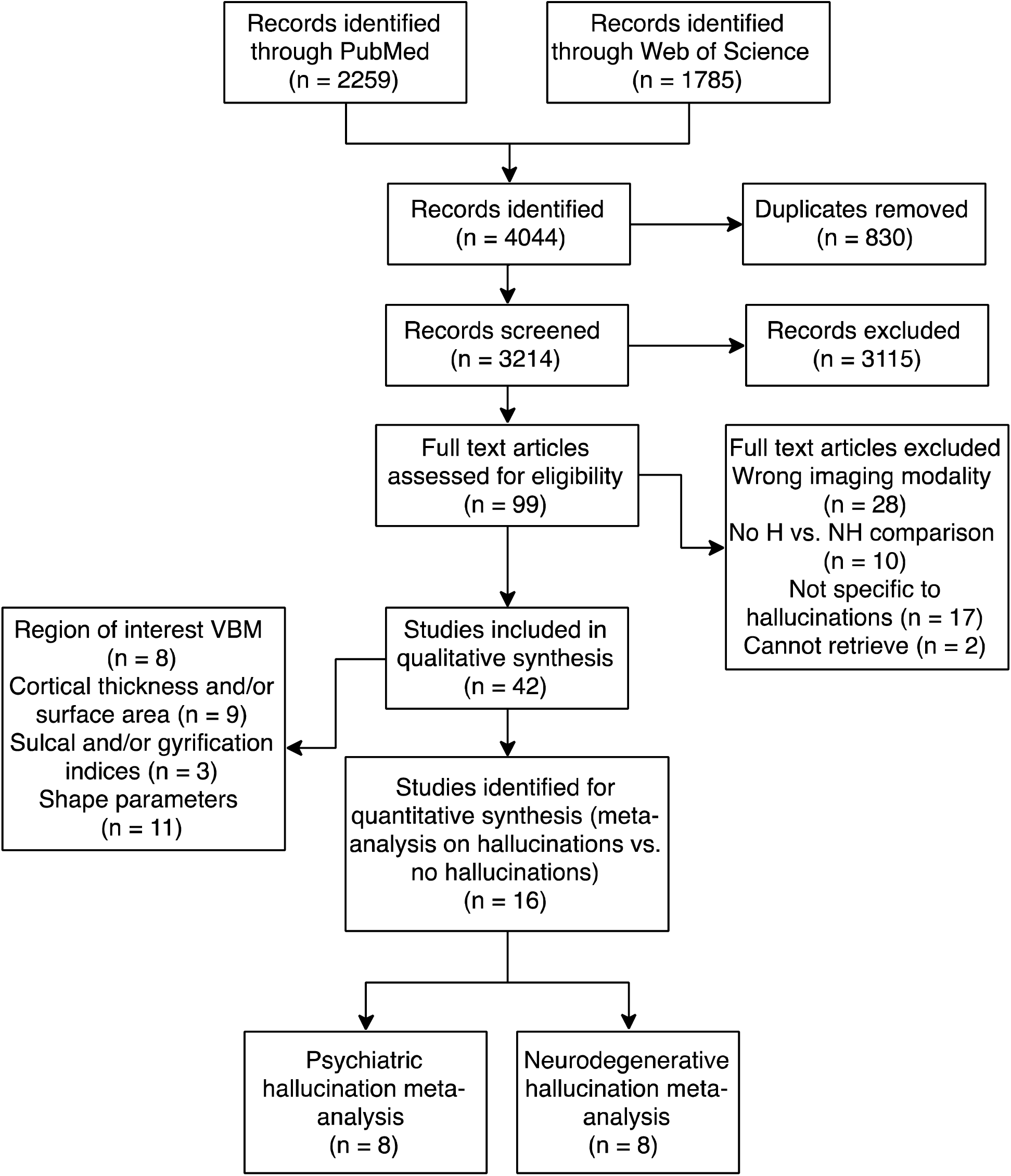
PRISMA flowchart for identification and selection of studies. Some studies performed analyses of multiple structural features and are therefore represented more than once. Abbreviations: H: population with hallucinations; NH: population without hallucination; VBM: voxel-based morphometry

In patients with hallucinations, relative to those without, GM reductions were identified in the left insula, right inferior frontal gyrus (IFG), left anterior cingulate/paracingulate gyrus, and left middle temporal gyrus (MTG), while GM increases were observed in bilateral fusiform gyrus (Table 3, Figure 3). Significant decreases in GM were apparent in six brain regions in patients with hallucinations compared to those without: (1) left lingual gyrus; (2) right supramarginal gyrus / parietal operculum; (3) left fusiform gyrus; (4) left parahippocampal gyrus; (5) right thalamus; (6) right lateral occipital gyrus (Table 3, Figure 3). Individuals with psychiatric relative to neurodegenerative hallucinations showed decreased GM in the left insula and anterior cingulate/paracingulate gyrus, and greater GM in the right lingual gyrus, IFG, and supramarginal gyrus, left thalamus, fusiform gyrus, inferior occipital gyrus, parahippocamapal and hippocampal gyri, and bilateral SFG (Table 4, Figure 3).

**Table 3.** Regions of significant differences in gray matter between patients with hallucinations compared to those without for psychiatric and neurodegenerative disorders.

| Group | Contrast | Region | Peak local maximum |  |  |  | Jackknife sensitivity analysis* |
| --- | --- | --- | --- | --- | --- | --- | --- |
|  |  |  | MNI coordinate | cluster size (no. of voxels) | SDM Z-score | uncorrected p-value |  |
| Psychiatric | H<NH | L insula | -46,2,-2 | 820 | -1.885 | 0.0000464 | 7/8 |
|  |  | R inferior frontal gyrus, pars triangularis / frontal pole | 48,36,8 | 281 | -1.464 | 0.0008257 | 7/8 |
|  |  | L anterior cingulate gyrus / paracingulate gyrus | 0,36,-2 | 132 | -1.259 | 0.0028023 | 7/8 |
|  |  | L middle temporal gyrus | -58,-42,-2 | 30 | -1.259 | 0.0028023 | 7/8 |
|  | H>NH | R fusiform gyrus | 44,-64,-18 | 574 | 1.455 | 0.0000877 | 7/8 |
|  |  | L lateral occipital cortex / fusiform gyrus | -40,-82,-16 | 345 | 1.454 | 0.0000981 | 7/8 |
| Neuro-degenerative | H<NH | L lingual gyrus / intracalcarine cortex | 0,-86,-4 | 1275 | -2.621 | 0.0000103 | 8/8 |
|  |  | L fusiform gyrus / inferior temporal gyrus | -36,-18,-26 | 50 | -1.860 | 0.0009702 | 7/8 |
|  |  | R supramarginal gyrus / parietal operculum | 54,-36,30 | 75 | -1.609 | 0.0034835 | 6/8 |
|  |  | L parahippocampal gyrus | -38,-32,-10 | 42 | -1.740 | 0.0018579 | 7/8 |
|  |  | R thalamus | 2,-2,12 | 14 | -1.637 | 0.0030603 | 7/8 |
|  |  | R lateral occipital cortex | 36,-80,14 | 10 | -1.511 | 0.0043970 | 6/8 |

Abbreviations: H: Hallucinations; NH: No hallucinations.
\*The jackknife sensitivity analysis tests the reproducibility of significant brain regions by iteratively repeating the statistical analysis, but systematically excluding one study from each replication<sup>55</sup>. Fractions show the number of study combinations in which the region was preserved out of the total number of dataset combinations.

**Table 4.** Regions of significant differences in gray matter between psychiatric and neurodegenerative hallucinations.

| Contrast | Region | Peak local maximum |  |  |  |
| --- | --- | --- | --- | --- | --- |
|  |  | MNI coordinate | cluster size (no. of voxels) | SDM Z-score | uncorrected p-value |
| Psychiatric < neurodegenerative | L insula | -42,-2,2 | 1784 | 1.794 | <0.0001 |
|  | L anterior cingulate gyrus / paracingulate gyrus | 0,44,-10 | 372 | 1.235 | 0.0011147 |
| Neurodegenerative < psychiatric | R lingual gyrus | 4,-84,-6 | 1080 | -2.331 | 0.0000206 |
|  | L superior frontal gyrus | -10,26,64 | 167 | -1.403 | 0.0016670 |
|  | R supramarginal gyrus | 52,-34,28 | 131 | -1.365 | 0.0020230 |
|  | L thalamus | -4,-4,10 | 115 | -1.516 | 0.0008154 |
|  | L fusiform gyrus | -24,-2,-42 | 90 | -1.494 | 0.0010064 |
|  | R inferior frontal gyrus, pars triangularis | 42,24,8 | 82 | -1.444 | 0.0013469 |
|  | L inferior occipital gyrus | -44,-78,-16 | 71 | -1.482 | 0.0010786 |
|  | L parahippocampal gyrus | -32,-18,-26 | 51 | -1.450 | 0.0013160 |
|  | R superior frontal gyrus | 14,36,-30 | 34 | -1.515 | 0.0008464 |
|  | L hippocampus | -36,-34,-8 | 33 | -1.524 | 0.0007690 |

**Figure 3.**
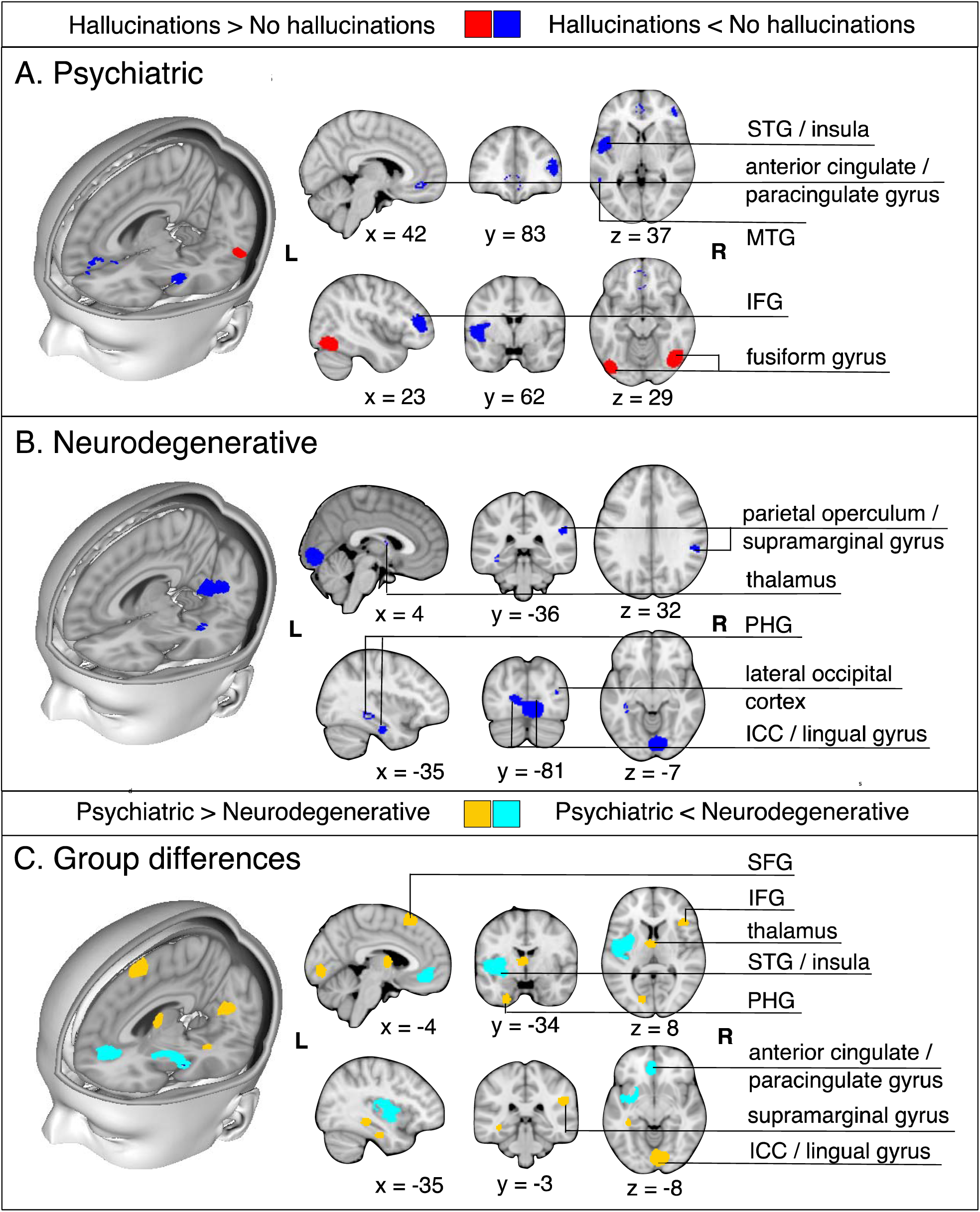
Meta-analysis results for individuals with hallucinations compared to those without hallucinations in psychiatric (A) and in neurodegenerative disorders (B). **A**. For psychiatric disorders, the meta-analysis revealed gray matter decreases in the left insula, right inferior frontal gyrus (pars triangularis) / frontal pole, left anterior cingulate gyrus / paracingulate gyrus, left middle temporal gyrus, and gray matter increases in the bilateral fusiform gyrus in patients with hallucinations relative to those without. **B**. For neurodegenerative disorders, the meta-analysis revealed decreases in the left lingual gyrus / intracalcarine cortex, left fusiform gyrus, right supramarginal gyrus, left parahippocampal gyrus, right thalamus, and right lateral occipital cortex. **C.** Formal comparison between meta-analyses revealed reduced GM in the left insula and left anterior cingulate/paracingulate gyrus for individuals with psychiatric relative to neurodegenerative hallucinations, and greater GM in the right lingual gyrus, IFG, and supramarginal gyrus, left thalamus, fusiform gyrus, inferior occipital gyrus, parahippocamapal and hippocampal gyri, and bilateral SFG. Abbreviations: STG: superior temporal gyrus; MTG: middle temporal gyrus; IFG: inferior frontal gyrus; PHG: parahippocampal gyrus; ICC: intracalcarine cortex; SFG: superior frontal gyrus

28 studies employed a regional and/or non-voxelwise approach to evaluate structural MRI data with respect to hallucination status: seven studies performed VBM restricted to predefined ROIs^43,61,69-73^, one performed source-based morphometry^74^, nine explored cortical thickness (CT) and/or surface area^75-83^, three investigated gyral/sulcal properties^43,84,85^, and 11 assessed structure-specific shape parameters^43,81,83,86-93^. Results are summarized in Tables 5–6. Overall, findings were heterogeneous, with few direct replications. In schizophrenia, the most consistent findings were reductions in CT in the left or right temporal gyrus for patients with hallucinations compared to those without^75,76,94^, coincident with the reductions in GM in left MTG observed in the meta-analysis (Figure 3). However, two studies reported increases in GM in temporal regions with hallucinations^88,90^. Hallucinations in PD and DLB were characterized by distributed patterns of cortical thinning^81,82^ and related to hippocampal volume, though the direction of this association was mixed^81,83^.

**Table 5.**
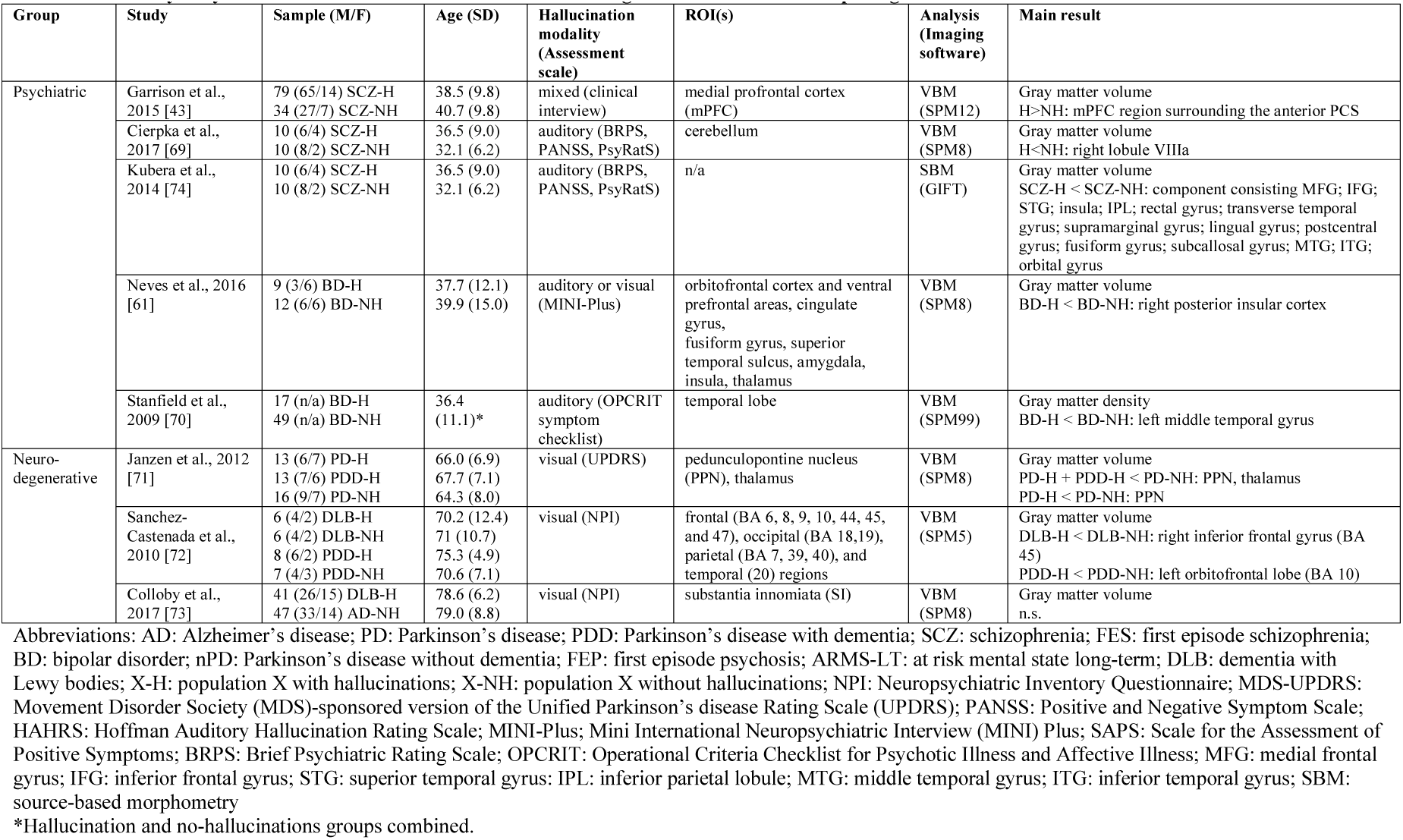
Summary of systematic review from GMV ROI studies of regional brain volume comparing individuals with and without hallucinations.

| Group | Study | Sample (M/F) | Age (SD) | Hallucination modality (Assessment scale) | ROI(s) | Analysis (Imaging software) | Main result |
| --- | --- | --- | --- | --- | --- | --- | --- |
| Psychiatric | Garrison et al., 2015 [43] | 79 (65/14) SCZ-H<br>34 (27/7) SCZ-NH | 38.5 (9.8)<br>40.7 (9.8) | mixed (clinical interview) | medial prefrontal cortex (mPFC) | VBM (SPM12) | Gray matter volume<br>H>NH: mPFC region surrounding the anterior PCS |
|  | Cierpka et al., 2017 [69] | 10 (6/4) SCZ-H<br>10 (8/2) SCZ-NH | 36.5 (9.0)<br>32.1 (6.2) | auditory (BRPS, PANSS, PsyRatS) | cerebellum | VBM (SPM8) | Gray matter volume<br>H<NH: right lobule VIIa |
|  | Kubera et al., 2014 [74] | 10 (6/4) SCZ-H<br>10 (8/2) SCZ-NH | 36.5 (9.0)<br>32.1 (6.2) | auditory (BRPS, PANSS, PsyRatS) | n/a | SBM (GIFT) | Gray matter volume<br>SCZ-H < SCZ-NH: component consisting MFG; IFG; STG; insula; IPL; rectal gyrus; transverse temporal gyrus; supramarginal gyrus; lingual gyrus; postcentral gyrus; fusiform gyrus; subcallosal gyrus; MTG; ITG; orbital gyrus |
|  | Neves et al., 2016 [61] | 9 (3/6) BD-H<br>12 (6/6) BD-NH | 37.7 (12.1)<br>39.9 (15.0) | auditory or visual (MINI-Plus) | orbitofrontal cortex and ventral prefrontal areas, cingulate gyrus, fusiform gyrus, superior temporal sulcus, amygdala, insula, thalamus | VBM (SPM8) | Gray matter volume<br>BD-H < BD-NH: right posterior insular cortex |
|  | Stanfield et al., 2009 [70] | 17 (n/a) BD-H<br>49 (n/a) BD-NH | 36.4 (11.1)* | auditory (OPCRIT symptom checklist) | temporal lobe | VBM (SPM99) | Gray matter density<br>BD-H < BD-NH: left middle temporal gyrus |
| Neuro-degenerative | Janzen et al., 2012 [71] | 13 (6/7) PD-H<br>13 (7/6) PDD-H<br>16 (9/7) PD-NH | 66.0 (6.9)<br>67.7 (7.1)<br>64.3 (8.0) | visual (UPDRS) | pedunclopontine nucleus (PPN), thalamus | VBM (SPM8) | Gray matter volume<br>PD-H + PDD-H < PD-NH: PPN, thalamus<br>PD-H < PD-NH: PPN |
|  | Sanchez-Castaneda et al., 2010 [72] | 6 (4/2) DLB-H<br>6 (4/2) DLB-NH<br>8 (6/2) PDD-H<br>7 (4/3) PDD-NH | 70.2 (12.4)<br>71 (10.7)<br>75.3 (4.9)<br>70.6 (7.1) | visual (NPI) | frontal (BA 6, 8, 9, 10, 44, 45, and 47), occipital (BA 18,19), parietal (BA 7, 39, 40), and temporal (20) regions | VBM (SPM5) | Gray matter volume<br>DLB-H < DLB-NH: right inferior frontal gyrus (BA 45)<br>PDD-H < PDD-NH: left orbitofrontal lobe (BA 10) |
|  | Colloby et al., 2017 [73] | 41 (26/15) DLB-H<br>47 (33/14) AD-NH | 78.6 (6.2)<br>79.0 (8.8) | visual (NPI) | substantia innomiata (SI) | VBM (SPM8) | Gray matter volume<br>n.s. |

Abbreviations: AD: Alzheimer's disease; PD: Parkinson's disease; PDD: Parkinson's disease with dementia; SCZ: schizophrenia; FES: first episode schizophrenia; BD: bipolar disorder; nPD: Parkinson's disease without dementia; FEP: first episode psychosis; ARMS-LT: at risk mental state long-term; DLB: dementia with Lewy bodies; X-H: population X with hallucinations; X-NH: population X without hallucinations; NPI: Neuropsychiatric Inventory Questionnaire; MDS-UPDRS: Movement Disorder Society (MDS)-sponsored version of the Unified Parkinson's disease Rating Scale (UPDRS); PANSS: Positive and Negative Symptom Scale; HAHRs: Hoffman Auditory Hallucination Rating Scale; MINI-Plus; Mini International Neuropsychiatric Interview (MINI) Plus; SAPS: Scale for the Assessment of Positive Symptoms; BRPS: Brief Psychiatric Rating Scale; OPCRIT: Operational Criteria Checklist for Psychotic Illness and Affective Illness; MFG: medial frontal gyrus; IFG: inferior frontal gyrus; STG: superior temporal gyrus; IPL: inferior parietal lobule; MTG: middle temporal gyrus; ITG: inferior temporal gyrus; SBM: source-based morphometry
\*Hallucination and no-hallucinations groups combined.

**Table 6.**
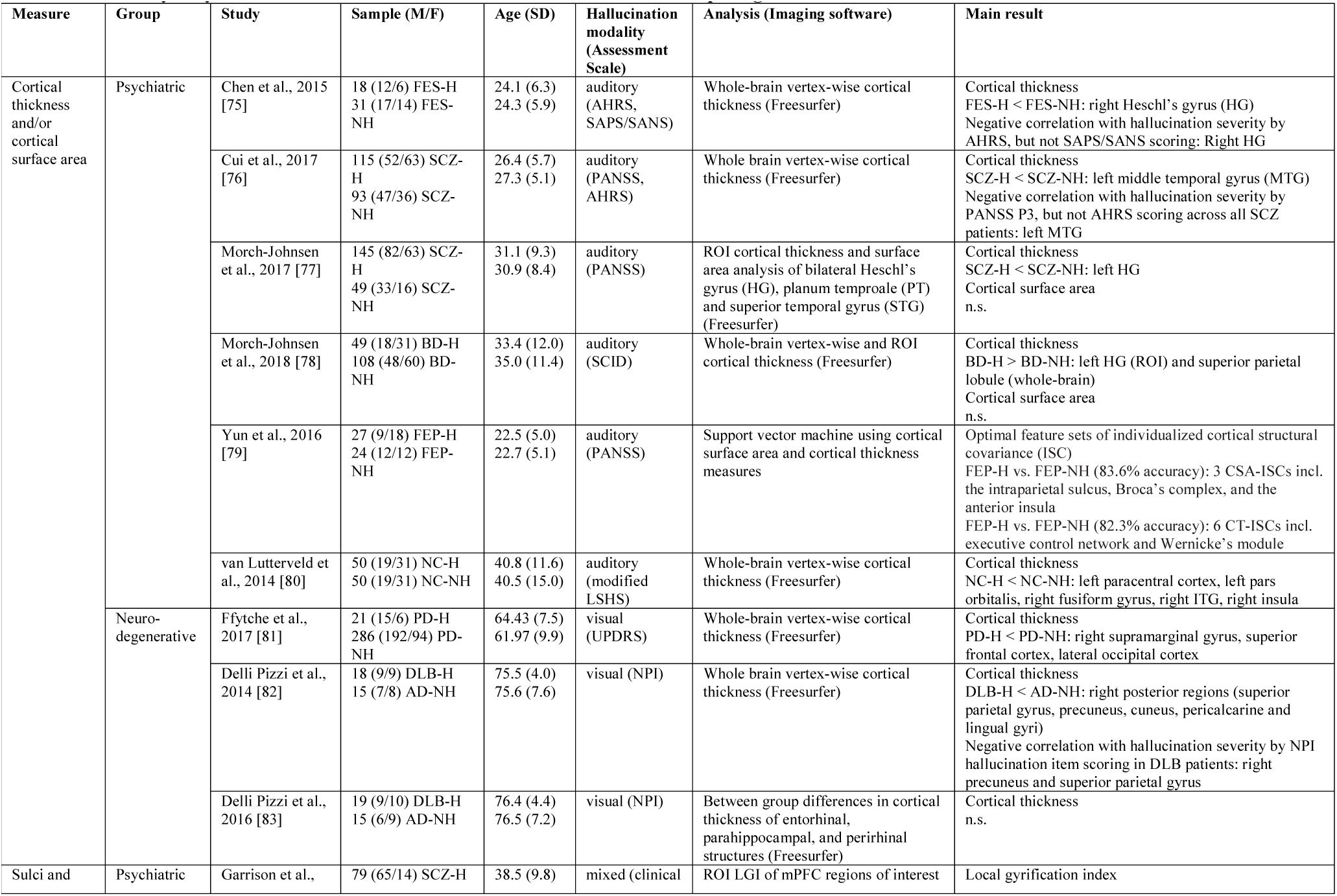
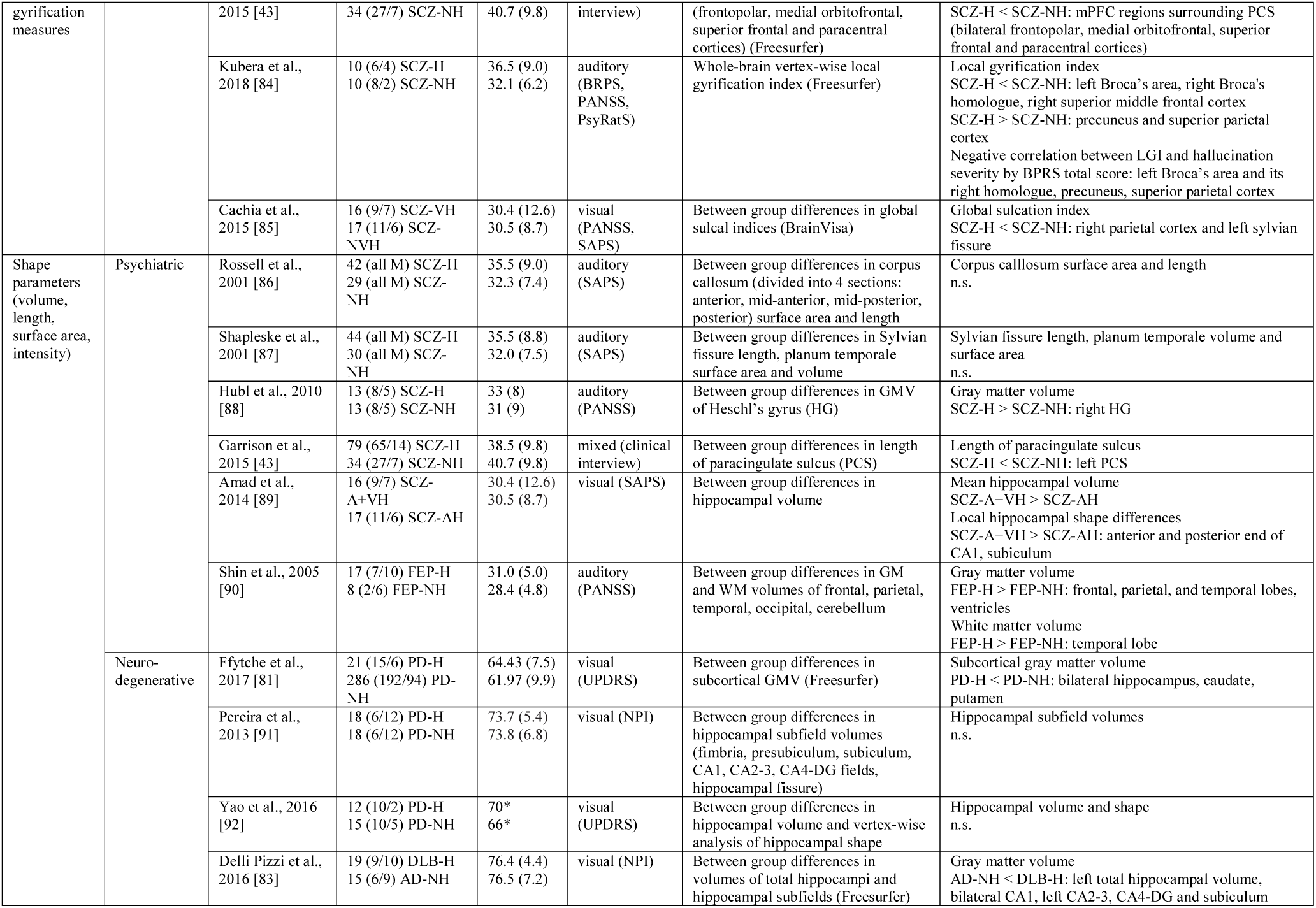
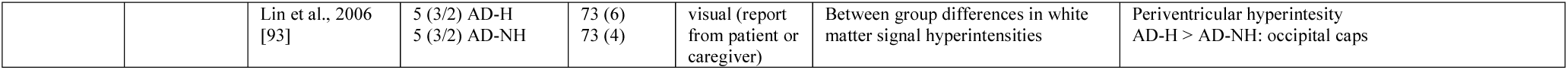
Summary of systematic review from non-voxelwise structural studies comparing individuals with and without hallucinations.

## Discussion

Distinctive patterns of neuroanatomical alteration characterize hallucination status in patients with psychiatric and neurodegenerative diseases, with the former associated with fronto-temporal deficits and the latter with medial temporal, thalamic and occipital deficits. These results broadly align with prior meta-analyses investigating GM correlates of hallucination severity of AVH in schizophrenia^23,24,26^ (Supplementary S3–4) and qualitative reviews on structural imaging studies of visual hallucinations (VH) in neurodegenerative illnesses^25,96^. The distributed pattern of structural changes seen in both hallucination signatures is suggestive of impairment in the coordination of information flow. Indeed, AVH in schizophrenia has been associated with increased functional activation in the STG, insula, anterior cingulate, and pre/post central gyrus^21,22^, reduced resting connectivity between default mode regions^97^, disruptions to the salience network^30^, and altered interactions between resting-state networks^97^. VH in PD have been associated with increased functional activity in the lingual gyrus, cuneus, and fusiform gyrus^27^, and hyperconnectivity in the default mode network^98^. Cortical thickness studies lend further support for divergent structural patterns, showing localized decreases in CT in temporal regions in schizophrenia spectrum disorders and more widespread decreases in dementia and PD.

We reviewed the brain structural abnormalities associated with hallucinations, yet how changes to the brain’s topological substrate translate to changes in an individual’s experiential landscape remain unknown. Our findings are consistent with multiple models of hallucinations (Figure 1). For instance, volume loss in temporal regions could reflect the misattribution of inner speech to a non-self source (inner speech model)^45^, or relate to abnormalities in cortical feedback for predictive signal processing (predictive processing account)^99^, or could be the result (or cause) of heightened resting state activity in the auditory cortex (resting state hypothesis)^29^, or a combination of some or all of these mechanisms. That substantial heterogeneity was observed in ROI VBM hypothesis-driven studies further emphasizes the limits of current theories.

Our meta-analyses suggest that there are at least two broad biological categories of hallucination mechanism: a psychiatric mechanism and a neurodegenerative mechanism. In support, structural signatures of hallucinations in the psychiatric meta-analysis overlap with comparisons of patients to non-disordered controls. For instance, a meta-analysis of GM changes in patients with psychosis compared to healthy controls shows reductions in bilateral insula and anterior cingulate cortex^100^, coinciding with regions identified in the meta-analysis of hallucinations in neurodevelopmental disorders, while thalamic, hippocampal, and occipital GM reductions in PD^101^ partly coincide with the changes seen in neurodegenerative hallucinations. The relation between disorder-specific GM changes and hallucination category suggests that hallucinations share networks of brain regions with the pathologies of the disorder in which they are embedded.

Knowledge of the structural correlates of hallucination types may help understand their cognitive phenotypes. For instance, hallucinations are linked to reality monitoring, the cognitive capacity to distinguish between self-generated and external sources of information^102^. Impaired in schizophrenia, reality monitoring is associated with the structure and function of the anterior cingulate cortex^43,102^. The cingulate gyrus is part of a network involving the IFG, ventral striatum, auditory cortex, right posterior temporal lobe whose functional connectivity is related to the subjective extent to which a hallucination feels real ^103^. Indeed, we propose that connectivity is key: together with the insula, the anterior cingulate constitutes nodes of the salience network, dysfunctions in which have been proposed as central to experiencing hallucinations^30^. Structural deficits in the insula in psychosis might also underpin atypical interactions between the DMN and salience network observed in hallucinations ^39^. The left STG/MTG have been robustly implicated in the manifestation of AVH^23,99^, emphasizing the importance of speech perception and processing in hallucinations in schizophrenia spectrum psychosis.

If hallucinations experienced by those with schizophrenia spectrum and bipolar psychosis are an example of a broader mechanism, then we predict that other neurodevelopmental disorders will have similar patterns of associated GM loss. For example, hallucinations have a prevalence of 43% in personality disorder^104^, suggested to be a neurodevelopmental disorder^105^, and are predicted to have a mechanism similar to other psychiatric disorders.

Abnormalities in the occipital cortex in neurodegenerative diseases suggest that deficits in sensory regions contribute to hallucinations of the associated sensory modality since VH are more common in PD than in schizophrenia^19^. Hallucinations in PD and AD were characterized by GM reduction in the thalamus and PHG. The thalamus relays information to higher level processing areas and contributes to working memory maintenance^106^, while the PHG is implicated in processing contextual associations in the service of memory formation and generating expectations about spatial relations^107^. Their involvement supports memory-related processes in hallucinations^28^, though may equally relate to neurodegenerative pathologies. The anterior cingulate was implicated in hallucinations occurring in psychiatric disorders, but not neurodegenerative aetiology. As the anterior cingulate is involved in self-referential processing, this is consistent with the observation that psychotic hallucinations address the individual and vary across continental location and historical time period^14,108^. Conversely, hallucinations in PD have a more passive quality and form historically stable categories of visual percepts^4^.

Anatomic heterogeneity related to hallucination presence/absence has important consequences for the plurality of treatment options. A specific example is repetitive transcranial magnetic stimulation (rTMS) used to reduce hallucination frequency and severity in schizophrenia, albeit with some reservations^109,110^. A number of parameters including frequency of stimulation and anatomical site contribute to the outcome of rTMS, and so anatomical heterogeneity is a possible source for the ambiguity of efficacy in therapeutic trials^111^. In general, effective treatment for hallucinations requires an understanding of the underlying mechanism that we suggest varies across diagnosis.

The multimodality of hallucinations is under-documented and under-researched, with <2% of studies included in this review probing hallucinations beyond audition or vision^64^. However, 30-50% of schizophrenia or PD patients report hallucinations in more than one modality^2,112^: olfactory hallucinations are present in 10-13.7%^51,113^ and tactile sensations frequently co-occur with auditory hallucinations^1^. Despite the dimensionality of hallucinations, many questionnaires and theoretical models target unimodal accounts. Non-clinical individuals who hallucinate or hear voices are receiving increasing interest in scientific research^7^, yet only one study in this review assessed a structural correlate (cortical thickness) of hallucinations in this population^80^.

The prevalence of hallucinations in the general population varies across the lifespan with peaks in early life (<30 years) and between 50-59 years^114^. Results from these meta-analyses predict that early onset of hallucinations will have a pattern of frontotemporal structural deficits similar to psychiatric disorders with neurodevelopmental origins, whilst later onset will show a neurodegenerative pattern of GM change in the occipital cortex, medial temporal lobe and thalamus. In any case, empirical neuroimaging and cognitive research in non-clinical groups and non-dominant modalities is necessary to extend the limits of current knowledge.

As with all meta-analyses, statistical power is restricted by the size of the extant literature, which in neuroimaging the experience of hallucinations remains immature. Despite this, the overall sample size was comparable to other SDM meta-analyses (n=233 H, n=194 NH for psychiatric; n=128 H, n=162 NH for neurodegenerative)^115,116^. The questionnaires used to assess hallucination status varied in the time frame bounding the hallucination, from within the current week to lifetime history, and few assessed phenomenological characteristics of hallucinations. The latter is potentially confounding as experiential differences may map to different neural substrates^117^. Understanding the neurobiology supporting the content of hallucinations may help in personalizing treatment strategies since hallucination content is related to cognitive profile in PD^118^. The divergence in our meta-analytic findings for psychiatric and neurodegenerative disorders may be partly attributable to differences in modality, since hallucinations experienced in schizophrenia spectrum and bipolar psychosis were predominantly auditory, while those in PD and AD were mostly visual. However, there was no overlap in brain regions identified in the two meta-analyses. Moreover, the reported modalities are partly construed by the questionnaire used. An important question is whether hallucinations of the same modality have common characteristics regardless of diagnosis. Finally, there was also a significant difference in the ages of the participants in the two meta-analyses, although each meta-analysis had its own age-matched control group and thus the comparison between disorders did not capture differences due to aging.

Hallucinations in clinical and non-clinical populations are diverse in content, modality, frequency, and affect, among other dimensions. Though hallucinations have been explored transdiagnostically at the level of phenomenology, little empirical work has made group comparisons of brain structure related to hallucinations. We show that hallucinations in psychiatric disorders have distinct neuroanatomical organization from the pattern observed in neurodegenerative diseases, and in doing so hypothesise at least two structural substrates associated with the hallucinatory experience. This categorical differentiation in the neurobiology of hallucinations is important for optimizing or developing treatment strategies, and makes specific predictions about other disorders, such as personality disorder, and the onset of hallucinations in the general population. The structural networks involved in hallucinations partly coincide with the respective case-control comparisons, and are thus embedded within the broader neuroanatomical phenotype, emphasising the importance of non-hallucinating patient control groups. Hallucinations are experienced in a variety of mental health contexts and are important phenomena in probing our perception of the external world, but theoretical work has not yet captured the diversity of hallucinations across modalities or diagnoses. By hypothesising at least two mechanisms for hallucinations, we suggest incorporating this plurality in future research. These meta-analyses offer a critical starting point.

## Contributors

JS conceived and directed the project. CR planned the search criteria, completed the literature search, data extraction, quality assessment, data analyses and summary, created the figures, and wrote the first draft of the manuscript, with input from JRG, JS, and GM. JRG, JS, and GM confirmed the results of data extraction. All authors critically reviewed the manuscript and contributed to its writing and revision.

## Declaration of interests

Ms. Rollins reports a scholarship from Gates Cambridge during the conduct of the study. Professor Rowe reports grants from Wellcome Trust during the conduct of the study, grants from NIHR, McDonnell Foundation, PSP Association, Parkinsons UK, Medical Research Council, Evelyn Trust, and AZ-Medimmune, personal fees from Asceneuron, and other from Guarantors of Brain outside the submitted work. Professor Suckling, Dr. Murray, Dr. Garrison, Dr. Simons, and Dr. O’Callaghan have nothing to disclose.

## Acknowledgements

The authors are supported by the following funding sources: Gates Cambridge (CR); Wellcome Trust (JBR, 103838; CO, 200181/Z/15/Z); National Health and Medical Research Council Neil Hamilton Fairley Fellowship (CO, 1091310); Wellcome Trust Collaborative award (JRG), and a joint award from the Medical Research Council and the Wellcome Trust (JSS).

